# Systematic annotation of hyper-variability hotspots in phage genomes and plasmids

**DOI:** 10.1101/2024.10.15.618418

**Authors:** Artyom A. Egorov, Vasili Hauryliuk, Gemma C. Atkinson

**Author notes:** To whom correspondence should be addressed: Artyom A. Egorov, Gemma C. Atkinson.

## Abstract

Bacterial and bacteriophage genomes contain genomic regions of hyper-variability (diversity hotspots) caused by insertions of mobile genetic elements (MGEs), non-homologous recombination events and non-horizontal hypermutation. Accessory genes encoded in the diversity hotspots are involved in anti-MGE defence and counter-defence, virulence and antimicrobial resistance (AMR), thus playing key roles in interactions amongst phages, MGEs, bacteria and eukaryotic hosts. To date the majority of research has been focused on either individual hotspots or on relatively limited sets of hotspots in a small set of genomes, typically from a single species. A global understanding of hotspot diversity and dynamics still lacking. To address this gap, we developed iLund4u, an algorithm for the systematic annotation of hotspots across millions of sequences. Using a proteome composition approach, iLund4u detects proteome communities, annotates accessory proteins and identifies hotspots. By analysing 873K phage genomes and 696K plasmid sequences we identified 13.7K hotspots and 171K diverse protein families encoded there as cargo. Furthermore, iLund4u allows for protein search and proteome annotation functions versus a precomputed iLund4u database. In the protein search mode iLund4u identifies all hotspots that encode homologues of a query protein. In the proteome annotation mode iLund4u annotates hotspots by searching for communities of similar proteomes. Detailed documentation, user guide and the source code are available at the iLund4u home page: art-egorov.github.io/ilund4u.

## Introduction

Microbial genomes are highly diverse, being shaped by various genetic variation-generating molecular mechanisms (Brockhurst *et al*, 2019; Kirchberger *et al*, 2020; Koonin & Wolf, 2010; Ryall *et al*, 2012). Horizontal gene transfer (HGT) plays a substantial role in this process and can result in the formation of discontinuous blocks of consecutive genes referred as genomic islands (GIs, or simply, islands) with differential presence and absence patterns across closely related genomes (Arnold *et al*, 2022; Juhas *et al*, 2009; Ochman *et al*, 2000). Islands are not evenly distributed along bacterial genomes. Rather, they are concentrated in hypervariable chromosomal regions (hotspots) characterised with rapid gene turnover and flanked by less variable regions (Lescat *et al*, 2009; Molari *et al*, 2024; Oliveira *et al*, 2017; Touchon *et al*, 2009). Mobile genetic elements (MGEs) such as plasmids and phages possess their own hotspots of variability that are shaped by recombination, transposons and gene swapping (Beamud *et al*, 2024; Weisberg & Chang, 2023; Yutin *et al*, 2024). Phage-encoded hotspots can contain so-called “moron” regions (a contraction of “more genes on”) defined as autonomously expressed modules of accessory genes (Cumby *et al*, 2012; Juhala *et al*, 2000; Rousset *et al*, 2022). Diversity-generating retroelements (GDRs) can also generate local variability, through mutagenic reverse transcription, without transfer of exogenous genetic material (Dore *et al*, 2024; Guo *et al*, 2008). These different processes that shape microbial genomes share one important property: they result in relatively variable loci surrounded by more conserved (core) genes. Here, we refer to such a locus in a particular genome/sequence as a “variable island”, while a group of variable islands that share the same conserved genomic neighbourhood in multiple genomes/sequences we call a “hotspot” **(Fig. 1)**.

**Figure 1.**
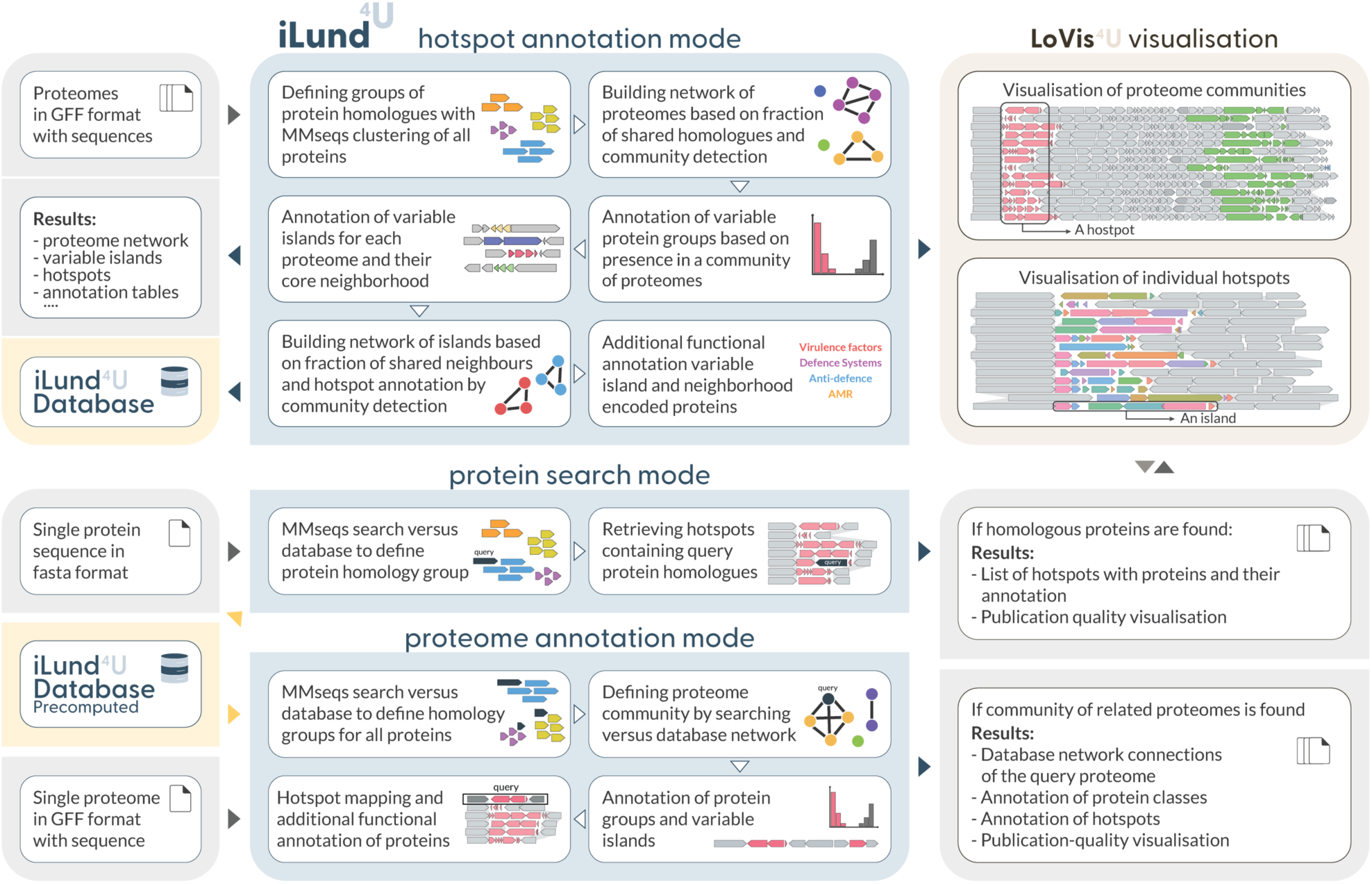
iLund4u workflow and functionality. Descriptions of the input data and the results structure (grey backdrop) and analysis steps (blue backdrop, *center*) are detailed for the hotspot annotation mode (*top*), protein search mode (*middle*) and proteome annotation mode (*bottom*). The LoVis4u (Egorov & Atkinson, 2024) library for genomic locus visualisation is illustrated with a schematic example of the output (brown backdrop, *top right*). The iLund4u database which is generated with the hotspot annotation mode serves as an input for protein search and proteome annotation modes (yellow backdrop, *left*).

Depending on the functions of their cargo genes, different islands have been assigned to categories as virulence, anti-microbial resistance (AMR), anti-MGE defence, anti-defence or pathogenicity (Dobrindt *et al*, 2004; Getz & Maxwell, 2024; Juhas *et al*., 2009; Makarova *et al*, 2011; Pinilla-Redondo *et al*, 2020; Schmidt & Hensel, 2004). This thematic functional clustering has been successfully exploited for the discovery of new biology. The tendency of anti-phage defence systems to co-localise in defence islands has enabled the “guilt by association” approach for the systematic discovery of new defence systems (Bernheim & Sorek, 2020; Doron *et al*, 2018). Analogously, the tendency of defence islands to be located in a particular hotspot position on chromosomes or MGEs, has been exploited for the discovery of new defence genes (Beamud *et al*., 2024; Hochhauser *et al*, 2023; Johnson *et al*, 2023; Koonin *et al*, 2017; Rousset *et al*., 2022). Finally, some hotspots have been reported to be enriched AMR, tailspike, or DNA polymerase cargo genes (Lescat *et al*., 2009; Pas *et al*, 2023; Yutin *et al*., 2024).

Current methods for annotation of hotspots tend to involve three approaches i) identification of loci within pre-defined conserved neighbourhoods, ii) association with known hotspot “cargo”, and iii) pan-genome methods. In an example of the first approach, conserved genes flanking hotspots in P2 phages and their P4 satellites enabled mining for new antiphage defence systems (Rousset *et al*., 2022). A combination of approaches has been used to identify defence hotspots in *Escherichia coli* and *Pseudomonas aeruginosa* pan-genomes, taking known defence systems as a start and then searching for conserved flanking gene arrangements (Hochhauser *et al*., 2023; Johnson *et al*., 2023). A similar approach based on known cargo has been used to annotate DNA polymerases and tailspike-containing hotspots in tailed phages (Pas *et al*., 2023; Yutin *et al*., 2024). A more systematic approach to annotation of new hotspots based on pan-genome analysis of regions of genome plasticity (RGPs) was implemented in panRGP (Bazin *et al*, 2020). This method uses a set of genomes of a given species to first define clusters of core, shell, and cloud genes (concepts reviewed in (Brockhurst *et al*., 2019; Eizenga *et al*, 2020)), and then annotates the RGPs that are flanked by conserved core genes. While this approach can be used on the host chromosomes of thousands of genomes of the same species, it cannot be readily extended to MGEs such as plasmids and phages, as they may have host ranges that extend beyond the species level, and have genome and population structures that are not suitable for taxonomy-based clustering. Additionally, an algorithm for detecting accessory genes and regions without prior genome clustering was implemented in a recent study (Silas *et al*, 2023). However, as this method relies on the analysis of pairwise gene combinations, scaling to large numbers of genomes is a challenge and limits systematic application.

To fill the gap in truly large-scale identification and annotation of hotspots across millions of sequences, we developed the iLund4u algorithm. Rather than being fixed to any formal taxonomic level, our approach first constructs a network of sequences based on proteome composition similarity. This allows us to apply iLund4u not only to sets of chromosomes or chromosome regions from different species but also to plasmid and viral sequences. After identifying and annotating communities of related sequences, iLund4u then searches for variable islands, and identifies hotspots based on island network partitions. We have applied the algorithm to 873 thousand (K) phage genomes retrieved from the PhageScope database (Wang *et al*, 2024) and 696K plasmid sequences from the Integrated Microbial Genomes & Microbiomes (IMG/M) database (Camargo *et al*, 2024). From these we identified 13.7K hotspots that encode cargo genes belonging to 171K diverse protein families, with the majority (∼89%) being functionally unannotated. We show that on average 9% of the length of each phage genome is comprised of hotspot islands, with a median value of 6.3%, and that the average total island length increases as the size of the phage genomes increase. The higher the diversity of island protein cargo functions, the longer the variable island. Leveraging these results, we have built a publicly available iLund4u database of phage and plasmid hotspots. Finally, in addition to hotspot annotation, iLund4u allows for user-input protein search and proteome annotation functions, taking advantage of the precomputed iLund4u database. In protein search mode, iLund4u identifies homologues of query proteins that are encoded in hotspots, providing additional information for functional prediction through the principles of guilt by association, embedding, and co-localization. In proteome annotation mode, iLund4u annotates hotspots by identifying communities of similar proteomes in a database, extending the functional annotation of new genomes including phages and plasmids. The iLund4u home page can be found at art-egorov.github.io/ilund4u.

## Results

### iLund4u is a fast and scalable method for annotating variable hotspots

We have recently identified several defence systems that are often located on variable regions of temperate phages, such as CapRel, CmdTAC, and HigBAC, (Mets *et al*, 2024; Zhang *et al*, 2022), aided by our gene neighbourhood analysis too FlaGs (Saha *et al*). However, a method that can automatically identify and visualise these regions across large datasets has been lacking. To address this, we developed iLund4u, a scalable tool capable of processing millions of sequences to systematically detect variability hotspots.

iLund4u has three modes of usage: hotspot annotation, protein search, and proteome annotation (**Fig. 1**). Hotspot annotation is the primary mode and is designed for identifying variability hotspots with protein-coding gene cargo in a given set of genomes. This mode includes several intermediate analysis steps, such as building a network of genomes based on proteome composition similarity, as well as subsequent clustering.

A key step in the iLund4u pipeline is construction of the proteome network and identification of communities of sequences that have similar proteome compositions. In short, at this step the input genomes are clustered based on the fraction of shared homologues without using taxonomical information or nucleotide sequence alignments. Similar clustering approaches based on proteome equivalence have been implemented in PhamClust (Gauthier & Hatfull, 2023) and in the “weighted gene repertoire relatedness” approach (Bobay *et al*, 2013; Pfeifer *et al*, 2021). However, unlike iLund4u, PhamClust uses hierarchical clustering and pairwise matrix which restricts scalability to sets containing more than thousands of sequences. Weighted gene repertoire relatedness, in turn, requires the processing of all-versus-all protein alignments, also limiting its application.

Defining clusters of similar proteomes allows the binning of proteins similarly to what is done in pan-genome analyses, assigning proteins to “*conserved”, “intermediate”* and “*variable”* groups that are equivalent to “*core”, “shell”,* and “*cloud”* terms (Brockhurst *et al*., 2019). After protein group assignment, iLund4u annotates variable islands, and identifies hotspots by merging islands with similar conserved neighbourhoods.

The database that is generated with the hotspot annotation mode then can be used for protein search and proteome annotation modes. iLund4u is distributed with a precomputed database for this, built on our analysis of hotspots in 873K phage and 696K plasmid sequences as described below. In the protein annotation mode, a user can determine whether homologues of a query protein are encoded in hotspots of the iLund4u database either as cargo or conserved flanking genes. The proteome annotation mode allows annotation of hotspots by finding communities of similar proteomes in the iLund4u database. Importantly, this mode can be run using a single proteome as input.

A detailed description of the workflow (also found in **Supplementary Text S1**) is available along with a parameter description at art-egorov.github.io/ilund4u.

### Phage genomes encode thousands of largely unexplored hotspots

To systematically annotate hotspots and build the iLund4u database for protein search and proteome annotation modes, we applied iLund4u to available phage sequences aggregated in the PhageScope database (Wang *et al*., 2024) (873K sequences as of version of September 2024) (**Fig. 2A**). As a pre-analysis step, we reannotated all phage nucleotide sequences using Pharokka (Bouras *et al*, 2023). Using the Pharokka-predicted protein sequences, we identified 275K proteome communities with the largest containing 1719 phages (**Fig. 2B**). 5.2K communities with size ≥10 were taken for the subsequent analysis steps. The distribution of the fraction of proteomes into “*variable*”, “*intermediate*”, and “*conserved* categories is U-shaped and follows the typical distribution seen in bacterial pan-genome analyses (**Fig. 2C**). (Domingo-Sananes & McInerney, 2021). The majority of protein groups (93.2%) are either “conserved” (17.5%) or “variable” (75.7%) with only 6.8% being “intermediate”. We then annotated 598K variable islands (3.1 on average per proteome), which were clustered into 12.4K hotspots (2.6 per community). The average number of island cargo proteins is three proteins, while the largest contains 116 proteins (**Fig. 2D**). Finally, 3.7K hotspots were merged to 1.7K non-singleton hotspot communities.

**Figure 2.**
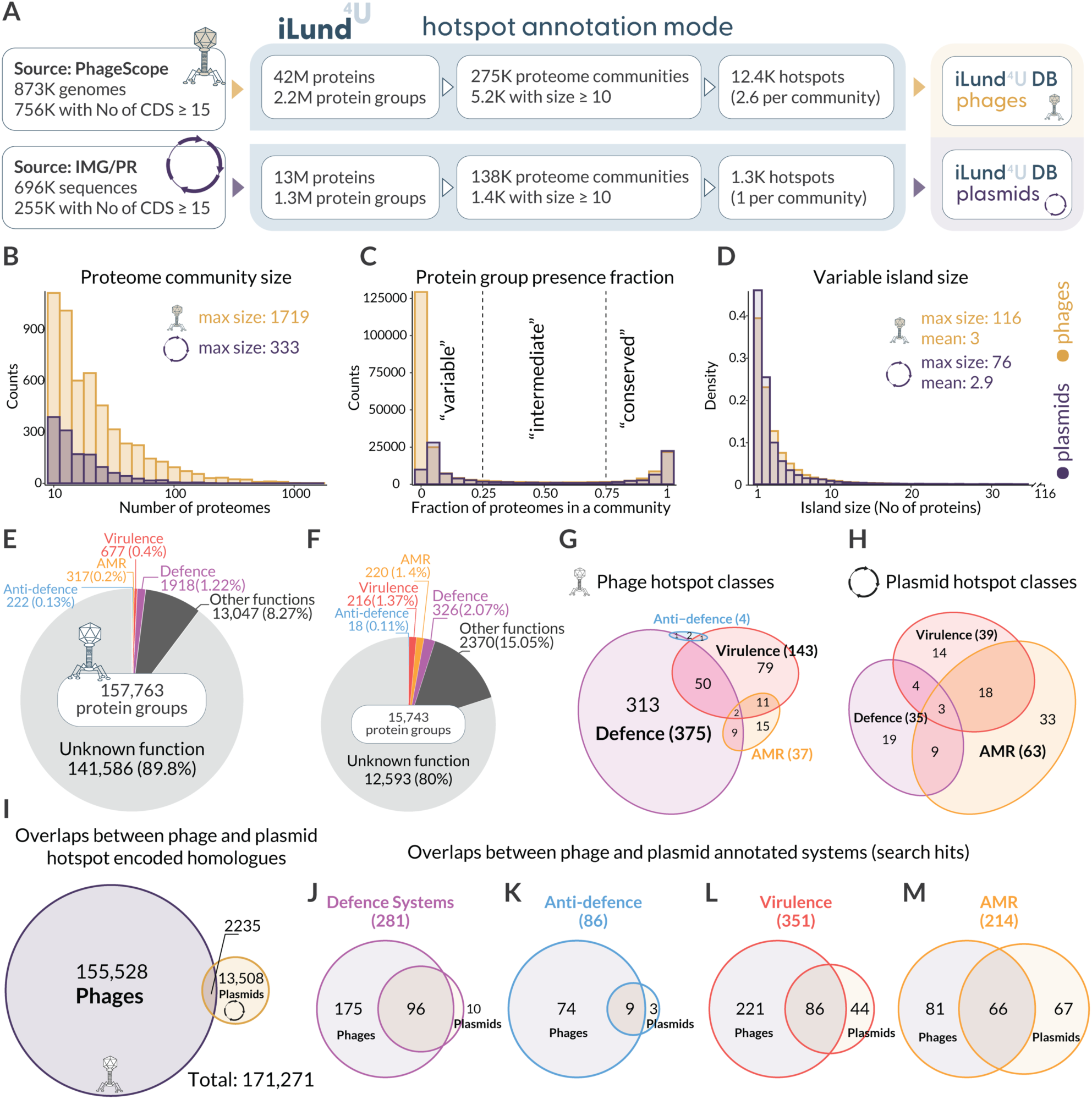
Discovery of thousands of hypervariability hotspots in phage genomes and plasmids. (**A**) The iLund4u workflow and summary statistics for hotspot annotation within phage and plasmid sequences. (**B**) Distribution of phage (yellow) and plasmid (purple) proteome community sizes (number of protein sequences). (**C**) Distribution of protein group fractions across proteome communities for phage (yellow) and plasmid (purple) protein variability groups. The fraction is the proportion of sequences within a proteome community that encode a particular group of protein homologues. Protein groups are classified as “conserved” if they are encoded in the majority of sequences (>75%). Those present in <25% sequences are considered “variable,” and the remainder are classified as “intermediate.” (**D**) Distribution of variable island sizes across hotspots, measured by the number of proteins. (**E-F**) Statistics of protein group classes according to the functional annotation for phages (E) and plasmid (F) proteins. (**G-H**) Statistics of the hotspot functional predictions for phage (G) and plasmid (H) hotspots. A particular function (e.g. defence) is attributed to the hotspot if at least 3% of its protein cargo groups has been annotated as homologues of the database member. Only hotspots with at least 50 cargo protein groups in total are considered. **(I)** Overlaps between phage and plasmid hotspot encoded homologues calculated based on the alignment of protein cluster representative sequences. **(J-M)** Overlaps of annotated function between phage and plasmid hotspot protein groups**: (J)** homologues of DefenseFinder (Couvin *et al*, 2018; Tesson *et al*, 2024) proteins that are encoded in phage or plasmid hotspots. **(K)** Proteins homologous to anti-defence proteins of dbAPIS (Yan *et al*, 2024). **(L)** Homologues of virulence factors in the VFDB (Liu *et al*, 2022) database. **(M)** AMRFinder (Feldgarden *et al*, 2021) database proteins.

The majority of hotspot-encoded homologous protein groups are functionally unannotated (141.5K out of 157.8K, 89.8%) (**Fig. 2E**). This is likely a large reservoir of variable accessory proteins that provide condition-specific fitness advantages (Brockhurst *et al*., 2019). Using functional annotation, we attempted to establish whether hotspots tend to have distinct functional “themes”. We applied the following rules: the hotspot should encode at least 50 diverse cargo protein groups with at least 3% of these groups having hits to the corresponding database (defence, anti-defence, AMR, virulence). In total we found 1327 hotspot communities with at least 50 cargo protein groups, 483 with at least one function being attributed and 74 of them (5.5% from the total) with more than one function (**Fig. 2G**).

### Plasmid-encoded hotspots are compositionally distinct from phage-encoded hotspots

Plasmids are known to carry accessory genes that affect bacterial phenotypes, and their repertoire has been shown to differ from those encoded by prophages (Takeuchi *et al*, 2023). To assess whether these genes are encoded in hotspots and to compare them with phage hotspots we applied the hotspot annotation mode to 696K plasmid sequences from the IMG/PR database (Camargo *et al*., 2024) (version of March 2024). We first excluded all (∼4K) sequences annotated as putative phage plasmids. (**Fig. 2A**). After the first filtering step, 255K sequences with proteome sizes of at least 15 proteins were classified into 138K proteome communities. 1.4K of these had at least 10 genomes and were taken for the subsequent analysis steps (**Fig. 2B**). As with phages, the fraction of proteomes in each variability group displays a U-shaped distribution with 88.7% being either “conserved” (35.9%) or “variable” (52.7%), and only being 11.3% annotated as “intermediate” (**Fig. 2C**). For 26.5K proteomes from the analysed communities, we identified 54K variable regions (2.04 on average per plasmid) that were clustered into 1279 hotspots (∼1 per community). The average number of cargo proteins encoded in each island is similar to that for phages, 2.9 (**Fig. 2D**). As with phages, the majority (80%) of plasmid hotspot-encoded cargo protein groups are unannotated. In the 137 plasmid hotspot communities containing at least 50 diverse protein cargo groups, 34 (24.8%) have more than one attributed function (**Fig 2H**).

We then evaluated the overlap between accessory genes encoded in phage and plasmid hotspots. To identify homologous groups, we aligned representative sequences from each cluster of phage homologues to those of plasmids using MMSeqs2 (Steinegger & Soding, 2017). Our analysis revealed 2235 shared protein groups and 171K distinct groups. Interestingly, while only 14.2% of all protein groups encoded in plasmid hotspots were also present in phage hotspots, the overlap was higher than on average for protein groups associated with defence, virulence, antimicrobial resistance (AMR), or anti-defence. For example, 91% of DefenseFinder system protein homologues identified in plasmid hotspots were also encoded in phage hotspots (**Fig. 2J**). Among the systems specific to plasmid hotspots, we identified the Avs3 system and several CRISPR-Cas related proteins. Avs3 binds to the monomer of the large terminase subunit and directly recognises the active site residues and ATP (Gao *et al*, 2020; Gao *et al*, 2022). Since the large terminase subunit is a signature protein for phages found in most genomes (Koonin *et al*, 2020), and Avs3 targets a wide variety of these proteins, the tendency for this system not to be encoded in phages may be explained by a risk of autoimmunity. Similarly, we analysed hotspot-localisation of anti-defence genes. Of the 86 anti-defence protein groups, nine are encoded in both phage and plasmid hotspots. This list includes six anti-CRISPR-Cas, two anti-restriction-modification systems (APIS036, APIS0063), and the anti-superinfection exclusion protein Cor (APIS032). An example of a specialised anti-defence phage hotspot is shown in **Supplementary Fig. S1**.

One of our plasmid communities containing the IncP-1beta multiresistance plasmid pB8 (AJ863570.1) corresponds to a set of conjugative plasmids that are transferred by from one bacterium to another via the pilus and have a specific directionality. The origin of transfer (*oriT*) of mobilizable elements defines the region on which the relaxosome complex is assembled and the region that will be transferred to the recipient cell first. This is called the leading region (Guasch *et al*, 2003). The transfer operon containing the relaxase gene is located near the *oriT* but is on the region that is transferred last. This is referred to as the lagging region of the plasmid (de la Cruz *et al*, 2010). The leading region can encode diverse anti-defence systems providing plasmid protection in upon conjugation (Samuel *et al*, 2024). Surprisingly, iLund4u identifies that the lagging strand can also carry hotspots. Specifically, we find a hotspot just downstream of the transfer operon with TraG, TraD, and TraC genes, containing AMR, and defence-associated genes (**Supplementary Fig. S2**). Across 88 diverse cargo protein groups encoded in this locus we detected 2 defence, and 15 AMR genes.

### Functionally related genes are organised in transcriptional units within variable islands of phage genomes

To understand the general features of phage-encoded islands, we analysed the diversity of variable islands in terms of size and composition. First, we computed the distributions of phage genome fractions that carry conserved genes and variable hotspots (**Fig. 3A**). On average approximately 9% of the phage genome is comprised of hotspot islands, with a median value of 6.3%. This agrees with a previous analysis of Mu-like phages that estimated that 6-10% of the genome is occupied by accessory genes (Cazares *et al*, 2014). The median value of the genome fraction occupied by regions encoding conserved genes is 74.1%. Note that even if a phage genome does not have a mappable hotspot, it can still encode variable and intermediate proteins in non-conserved locations, which explains the difference between the size of the conserved fraction and the size of the fraction that is not classified as hotspot.

**Figure 3.**
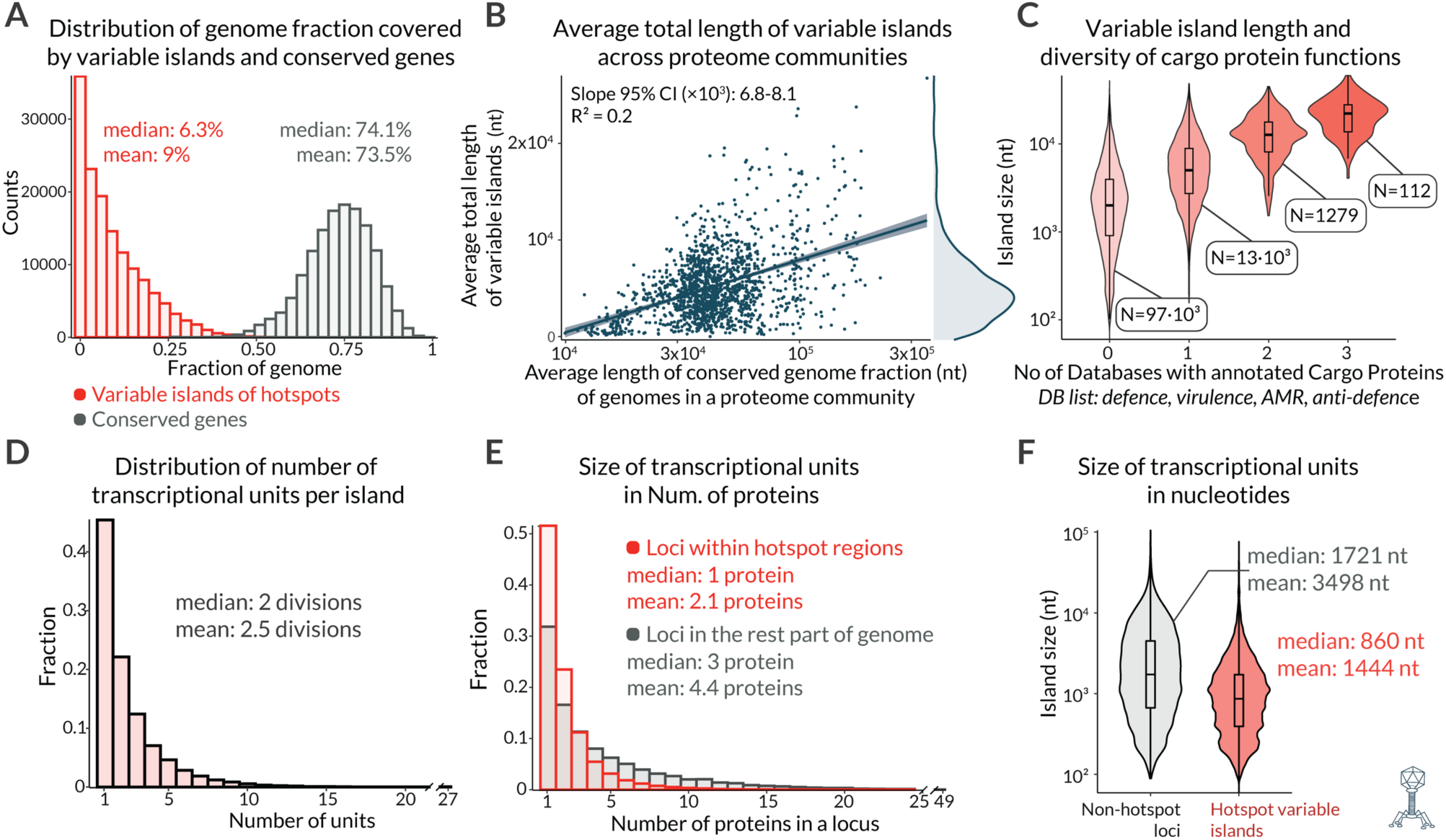
The architecture of phage variable islands. (**A**) Distribution of the phage genome fraction occupied by variable islands within hotspots (red) and conserved genes (grey). (**B**) Correlation between the average total length of variable islands across phages in a proteome community and the average length of the conserved genome fraction. (**C**) Distribution of variable island lengths based on the number of functional groups (including defence, virulence, AMR, and anti-defence) annotated within the island. (**D**) Distribution of the number of terminator sequence-based subdivisions (units) per variable island. (**E-F**) Comparison of the terminator sequence-defined transcriptional unit size, measured by the number of proteins (E) and nucleotides (F), for variable islands within hotspots (red) versus the rest of the genome (grey).

Next, we examined the relationship between hotspot length and phage genome length. We plotted the average total size of variable islands in nucleotides versus the average length of the conserved part of the genome within a community of phages in log scale and detected an expected positive correlation (95% CI of slope: 6.8-8.1 (x10^3^)) (**Fig. 3B**). We also found that larger hotspot islands (individually) were observed on larger phages (**Supplementary Fig. S3A**). There is a slightly negative correlation between the average fraction of the genome occupied by hotspot islands and the average length of the conserved fraction of the genome (**Supplementary Fig. S3B**). It is logical to assume that larger islands encode a greater functional diversity of their proteins. To test this, we compared the distribution of island size in nucleotides depending on the number of annotated functional groups for cargo proteins. Indeed, the median island size for an island having zero/one, two or three functional groups increases, with values 1374, 12364, and 22169 (nt), respectively (**Fig. 3C**).

Previous studies of phage morons have identified that they can consist of multiple transcriptional units (Cumby *et al*., 2012; Juhala *et al*., 2000). To identify such units in our phage hotspots, we predicted terminator sequences (Rho-dependent and Rho-independent) in all phage genomes using BATTER (Jin *et al*, 2023) and defined units as sets of adjacent genes with the same orientation divided by terminator sequences. We find there are on average 2.5 subdivisions (units) per island (**Fig. 3D**). When comparing unit size in hotspot islands versus unit size in the rest of the phage genomes, we find that hotspot units are usually smaller (2.1 versus 4.4 proteins on average (**Fig. 3E**), and nucleotide median length of 860 versus 1721 nucleotides (**Fig. 3F**). Finally, we evaluated the functional diversity of these units in comparison with the diversity of islands. While 1.24% of hotspot islands contain two or more functionally diverse cargo proteins, this value reduces to 0.1% for terminator-split units. It is important to note the caveat that almost in any random subdivision of islands we could expect a reduction in variability, and results are further complicated by the large fraction of unannotated genes. Nevertheless, our results indicate that guilt-by association prediction methods could be made more reliable by considering transcriptional units rather than whole islands. This is particularly likely to be the case if the island is large with many transcriptional units.

### iLund4u rediscovers well known P2-like phage hotspots

Some of the most well-studied phage-encoded hotspots are those found in P2-like phages. The classical P2 phage has two hotspots, one encoding the *old*-*tin* defence system (located between the nicking at origin of replication protein gpA and the portal protein) (Mosig *et al*, 1997; Myung & Calendar, 1995) and the other encoding *Z*/*fun* (located between tail-related proteins with a conserved baseplate protein upstream) (Calendar *et al*, 1998; Nilsson *et al*, 2004). These diversity regions have recently been extensively explored, yielding multiple new defence systems such as DP-T4-9 (CmdTAC), DP-T4-5, DP-T4-10, DP-α-5, and DprA-PRTase (Rousset *et al*., 2022; Vassallo *et al*, 2022). We asked whether iLund4u can rediscover these hotspots. Indeed, we identified and annotated hotspots in a representative subset of genomes from the P2-like phage community (15 out of 1008) (**Fig. 4A**). Both of the well-characterised P2 hotspots are identified (**Fig. 4B, C**).

**Figure 4.**
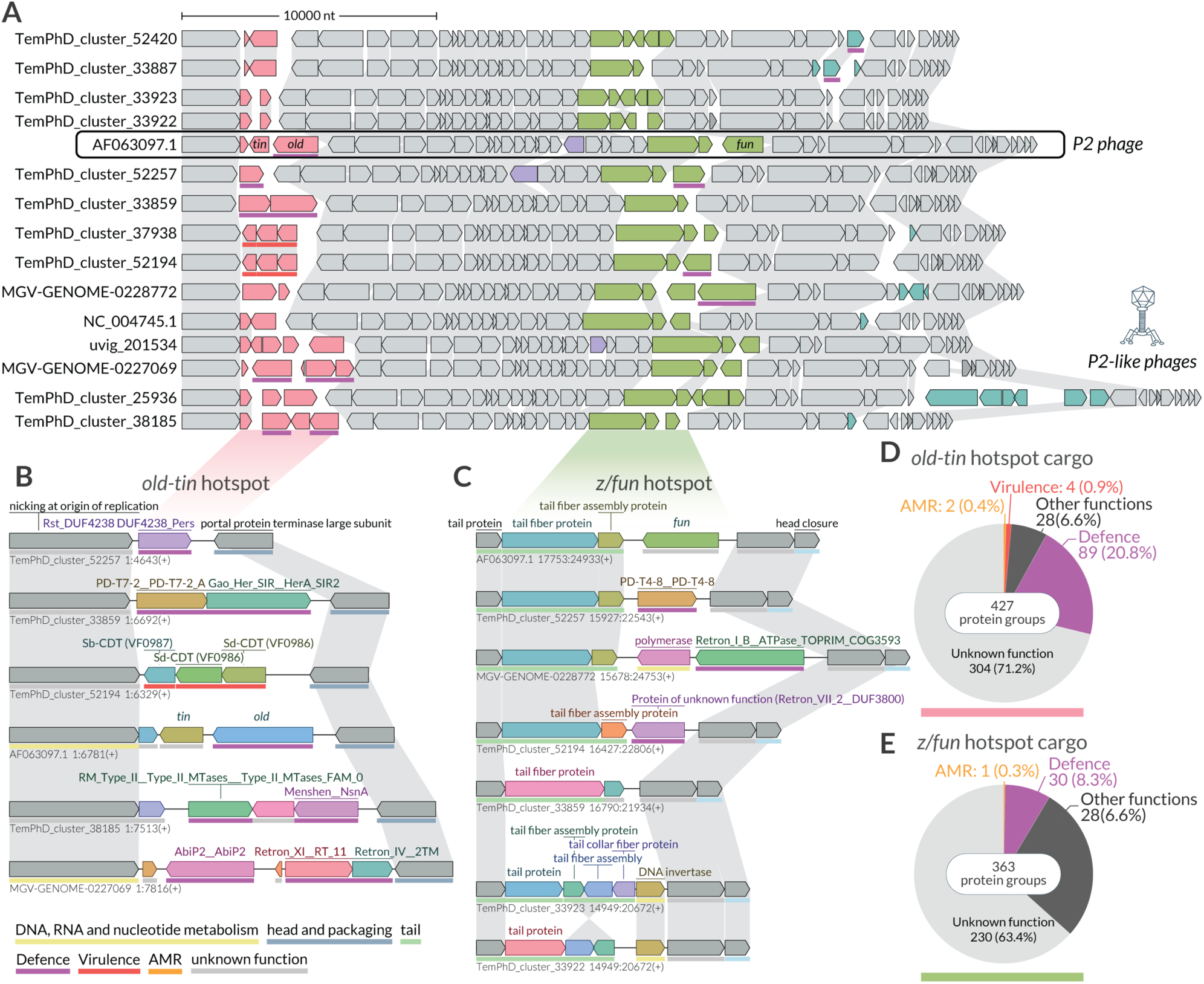
Hotspots of P2-like phages carry a diverse cargo of accessory genes. (**A**) Genomic comparison of selected P2-like phages. Conserved genes are shown in light grey, while both intermediate or variable protein-coding genes that reside outside of hotspots are dark grey. Hotspot cargo genes are highlighted in colour. (**B**) Detailed view of the *old*/*tin*-encoding hotspot, highlighted in red, which is located between the nicking at the origin of replication protein (upstream) and the portal protein (downstream). (**C**) Detailed view of the *Z*/*fun*-encoding hotspot, highlighted in green, situated downstream of the baseplate protein gene. Protein labels are hidden for hypothetical proteins or proteins of unknown function. Phage genomes were visualised using LoVis4u (Egorov & Atkinson, 2024), the functional annotation was made using PHROG (Terzian *et al*, 2021), and PyHMMER scan results are indicated by coloured lines beneath each locus, see the colour code at the bottom of the panel. **(D-E)** Pie charts of functional diversity of hotspot cargo protein groups.

Among the 427 diverse cargo protein groups of the *old*-*tin*-encoding hotspot, 89 (20.8%) have hits to the DefenseFinder database (Tesson *et al*., 2024). Other functional classes are also present. For example, as previously noted (Rousset *et al*., 2022), some phages encode in this hotspot a cytolethal distending toxin (CDT) (Hyma *et al*, 2005; Okuda *et al*, 1995), a toxic virulence factor that induces DNA damage and cell cycle arrest (Cortes-Bratti *et al*, 2001). In the *Z/fun*-encoding hotspot we also observed swapping events for tail fiber and tail fiber assembly genes, which encode receptor-binding proteins (RBPs) of phages and are responsible for host-range specificity (**Fig. 4C**). While such RBP modularity hotspots have been observed before (Pas *et al*., 2023), and we detect similar RBP regions in other phage communities, it is notable that this locus is mixed purpose, and can carry other categories of cargo, including other defence-like proteins (**Fig. 4C**). Additionally, we often see a DNA invertase encoded in this locus **(Fig. 4C)**, accompanied by the inversion of adjacent region genes. DNA invertases are particularly mobile and abundant in hotspots. For instance, one of the DNA invertase protein family was found on 108 diverse islands and 25 hotspots. iLund4u reveals a third hotspot of P2-like phages, located between a tail protein and the CI transcriptional regulator. Moreover, a hotspot in this locus has also been observed in other phage communities and previously reported in prophages (Rousset *et al*., 2022).

### A DGR containing hotspot in the human gut *Toutatisvirus* phages

iLund4u analysis of a cluster containing 173 *Faecalibacterium-*infecting *Toutatisvirus* phages reveals a highly diverse hotspot, both in terms of its size and composition **(Fig. 5)**. The hotspot is flanked by tail proteins upstream and an integrase downstream. The island size ranges from one to 23 protein coding genes, with 163 diverse cargo protein groups including AMR, Virulence, Defence, and components of diversity-generating retroelements (DGRs).

**Figure 5.**
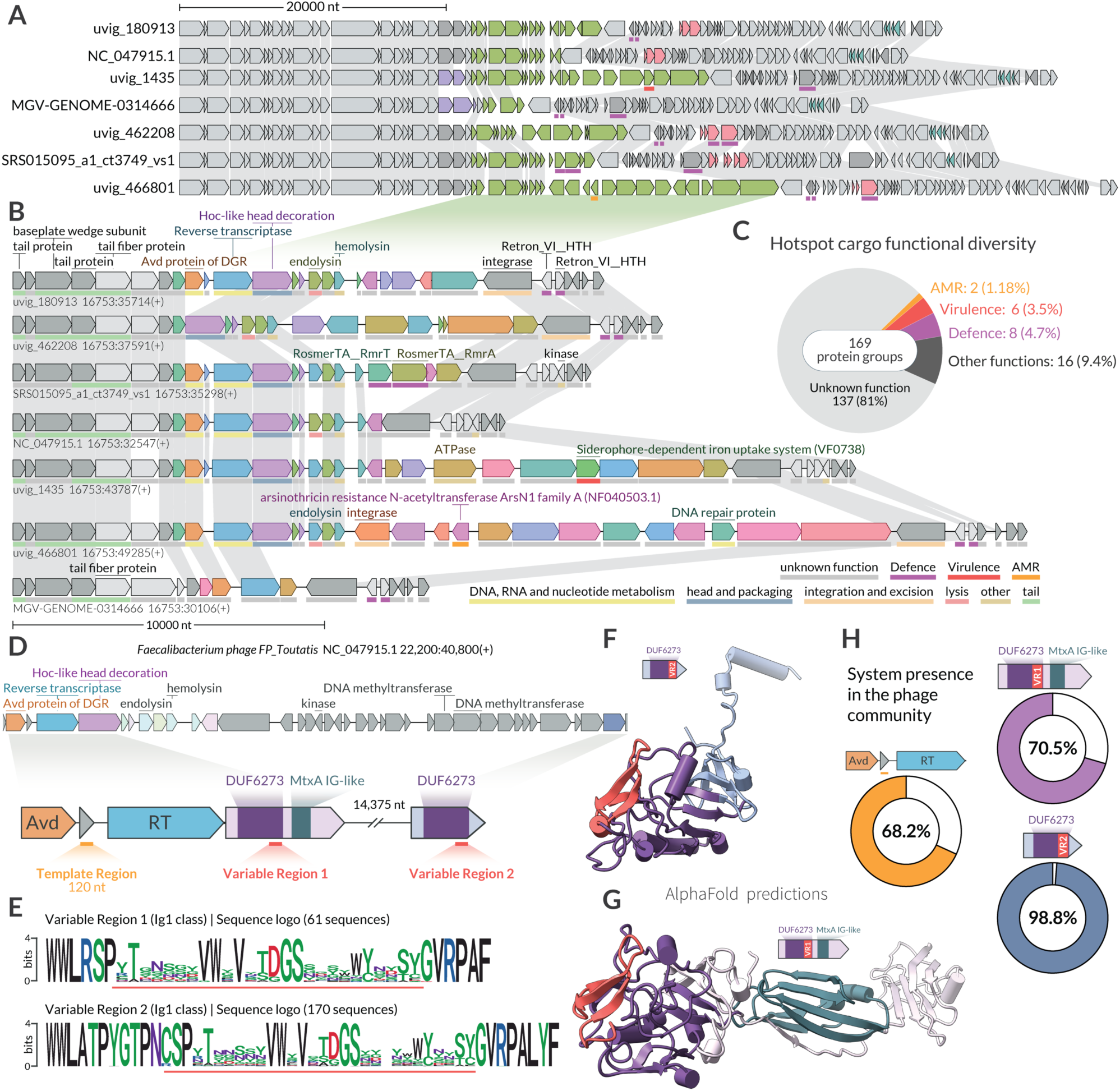
A mixed function hotspot of human gut phages shaped by a diversity-generating retroelement. (**A**) Genomic comparison of select *Toutatisvirus* phages. Conserved genes are shown in light grey, while both intermediate and variable protein-coding genes that reside outside of hotspots are dark grey. Hotspot cargo genes are highlighted in colour. (**B**) Detailed view of the DGR containing hotspot, highlighted in green, which is located between the tail proteins (upstream) and the integrase (downstream). Phage genomes were visualised using LoVis4u (Egorov & Atkinson, 2024), the functional annotation was made using PHROG (Terzian *et al*., 2021), and PyHMMER scan results are indicated by coloured lines beneath each locus, see the colour code at the bottom of the B panel. **(C)** Pie charts of functional diversity of hotspot cargo protein groups. **(D)** Detailed view of the DGR locus in the *Faecalibacterium phage FP_Toutatis* (NC_047915) with highlighted template and variable regions and domain organisation of the target genes. **(E)** Variable region (VR) sequence logos according to myDGR (Sharifi & Ye, 2019), which classifies both VRs as “Ig1 class”. **(F-G)** AlphaFold3 (Abramson *et al*, 2024) structure prediction of the target VR containing genes. **(H)** Presence statistics of the DGR system and the target genes in the phages of the community.

DGRs (Diversity-Generating Retroelements) are molecular systems that induce hypermutation in a variable region (VR) of the target gene, guided by the RNA encoded in a template region (TR) (Liu *et al*, 2002). In phages, DGRs frequently diversify receptor-binding proteins, such as tail fibers (Macadangdang *et al*, 2022). In this hotspot, DGRs are among the most abundant elements, being encoded in 118 (68.2%) of the phages within the community. Using myDGR (Sharifi & Ye, 2019), we annotated the DGR-associated TR and VR regions. The TR was located between the Avd and RT genes, as expected; however, instead of a single VR containing the target gene, we identified two distinct VRs **(Fig. 5D, E)**. The first VR-containing gene, annotated as Hoc-like head decoration, is also a hotspot member and is encoded downstream of the RT. The second VR-containing gene of unknown function is found 17,480 nt upstream of the TR and was part of the core genome, present in 98.8% of phages **(Fig. 5D, H).** Both genes contain a DUF6273 domain, with the VR located at its C-terminus. The VR1-containing gene also carries an MtxA Ig-like domain at the C terminus. Structural analysis reveals the DUF6273 domain resembles a C-type lectin (C-lec) domain **(Fig 5F, G)**. A similar locus with DGR targeting two DUF6273 domain-containing proteins was recently identified in gut phages (Baykov *et al*, 2023). The Ig-like domain-containing target is a member of the “tentaclin” (from TENTACLe + proteIN) family, while the second target resembles the Mtd-P1 protein, characterised by a C-lec-like domain and a VR at its C-terminus, functioning as a receptor-binding protein that interacts with pertactin (Miller *et al*, 2008). It is likely that these DUF6273 containing proteins function as receptor-binding proteins targeting either host surface or human intestinal cells. Curiously, in some phages, the DGR system is absent, yet both target genes remain present. It is possible that the main benefit of carrying DGR is as a rapid response diversity generator, in regions that have a selective pressure to diversify due to host interactions.

## Discussion

We introduce a novel method for the systematic annotation of genomic hyper-variability hotspots across millions of sequences. Analysis of hotspots and variable islands has previously led to the discovery of novel immune systems and counter-defence genes, relying on the “guilt-by-association” principle (Aravind, 2000; Galperin & Koonin, 2000; Koonin *et al*., 2017; Payne *et al*, 2024). Guilt by association takes advantage of the tendency for genes with similar functions to co-localise, forming functional islands (e.g. defence (Makarova *et al*., 2011), anti-defence (Pinilla-Redondo *et al*., 2020; Tesson *et al*., 2024). The mechanisms driving this co-localization are not fully understood, and might differ for MGE hotspots versus chromosomal islands (Koonin *et al*., 2017). To analyse the tendency for functional clustering within islands, we have evaluated thematic tendencies in our annotated phage islands. We find that islands do tend to have functional themes, as only 1,392 out of 111,979 islands (1.24%) encode proteins with more than one function across defence, virulence, AMR, and anti-defence. Notably, the median size of multi-functional islands is 12.9 MB, compared to just 2.2 MB for those with a single or no identifiable function. This suggests that larger islands should be treated more carefully when applying guilt by association for function prediction, and the information provided by iLund4u on island boundaries and sizes could be useful for selection of more functionally homogeneous islands. Furthermore, after subdividing these islands into transcriptional units, we find that only 0.1% remain multi-functional, indicating that the internal architecture of islands should also be taken into account when selecting hotspots of interest for biological discovery. We note, however, the caveat that the large number of proteins with unknown functions complicates more systematic analysis of functional specialisation, meaning that there is very likely some underestimation of the actual fraction of multi-functional islands.

The tendency for islands with specific functions to occupy the same hotspot positions can be formulated as a “guilt-by-location” approach (Beamud *et al*., 2024; Lescat *et al*., 2009; Rousset *et al*., 2022). We find thematic associations that suggest this is generally a useful strategy for functional prediction; among 375 hotspots with at least 50 cargo proteins and at least 3% annotated as defence, 83% (313) contained no other annotated functions. For other groups, the proportions of specialised hotspots were: 55% virulence, 41% AMR, and 50% anti-defence. While some hotspots do show clear functional specialisation, we observed greater diversity at the hotspot level (guilt by location) compared to individual islands (guilt by association). Functional specialisation likely depends on the constraints of transcriptional regulation, with anti-defence systems needing to be expressed early in the infection cycle, while defence systems are typically expressed during the lysogenic phase but not during the lytic phase.

We suggest the power of guilt-by-association and guilt-by-location approaches can be increased by combining them, lowering the risk of false predictions. The frequent horizontal transfer of accessory genes supports this, as 68.4% of defence-related protein groups were found in multiple islands, and over half (55.6%) appeared in more than one hotspot, with 9.4% in over or equal than ten. To facilitate this task of searching, we have developed the protein search mode of iLund4u that allows discovery and analysis of all phages or plasmids hotspots and islands where a query protein is encoded. We believe that our method and accompanying databases will be valuable for comparative genomic analysis and will facilitate functional annotation of accessory genes and particularly novel defence systems.

## Material and methods

### iLund4u implementation

iLund4u is written in Python3 and uses multiple python libraries: Biopython (Cock *et al*, 2009), bcbio-gff, scipy (Virtanen *et al*, 2020), configs, argparse, pandas (team, 2024), matplotlib (Hunter, 2007), seaborn (Waskom, 2021), progess, leidanalg (Traag *et al*, 2019), igraph (Csárdi & Nepusz, 2006), PyHMMER (Eddy, 2011; Larralde & Zeller, 2023), msa4u (Egorov & Atkinson, 2023) and lovis4u (Egorov & Atkinson, 2024). iLund4u also uses MMseqs2 (Steinegger & Soding, 2017) which is embedded in the library. MMseqs clustering and search parameters used are *“--cluster-mode 0 –cov-mode 0 -c 0.8 -s 7 --min-seq-id 0.25”* and *“-e 1e-5 -s 6”* respectively. For functional annotation, iLund4u uses the PyHMMER API (Eddy, 2011; Larralde & Zeller, 2023) (e-value cutoff 1e-3, query coverage cutoff 0.7, hmm coverage cutoff 0.5) searching a set of HMMs from the following databases: AMRFinderPlus (AMR genes) (Feldgarden *et al*., 2021), DefenseFinder and CasFinder (phage defence genes) (Couvin *et al*., 2018; Tesson *et al*., 2024), Cas dbAPIS_Acr (Anti-defence) (Yan *et al*., 2024), VFDB (Virulence factors) (Liu *et al*., 2022).

The python iLund4u package is available in PyPI (*python3 -m pip install ilund4u*), and the source code is provided on the GitHub page (github.com/art-egorov/ilund4u). Detailed documentation with an installation guide, and an example-driven manual are available at the iLund4u home page (art-egorov.github.io/ilund4u). A description of the pipeline steps is also available in the **Supplementary file Text S1**.

### Sources of phage and plasmid sequences

Phage sequences (873K) were downloaded from the PhageScope database (Wang *et al*., 2024) (version of September 2024) that aggerates sequences from multiple repositories: RefSeq (O’Leary *et al*, 2016), Genbank (Benson *et al*, 2018), EMBL (Kanz *et al*, 2005) and DDBJ (Ogasawara *et al*, 2020), and databases: PhagesDB (Russell & Hatfull, 2017), GOV2 (Gregory *et al*, 2019), GVD (Gregory *et al*, 2020), GPD (Camarillo-Guerrero *et al*, 2021), MGV (Nayfach *et al*, 2021), CHVD (Tisza & Buck, 2021), STV (Santos-Medellin *et al*, 2021), IGVD (Shah *et al*, 2023), IMG/VR (Camargo *et al*, 2023). As source of 700K plasmid sequences we used the IMG/PR database (Camargo *et al*., 2024) (version of March 2024). In addition, ∼4K plasmid sequences annotated as putative phage plasmids were excluded from the analysis.

### Data processing

Before using iLund4u we standardised annotation of phage and plasmid sequences. For phages we used Pharokka (Bouras *et al*., 2023) in ‘meta’ mode with the following optional arguments: “--meta --split --skip_mash --dnaapler”. For plasmid sequences we applied Prokka (Seemann, 2014) using the default parameters. Then gff files were used as input to iLund4u. We used iLund4u with two optional arguments: i) *-gct* option, which takes additional information on sequence circularity as input. ii) *-rnf* argument that forces to report also hotspots that flanked only from one side. Note, that all statistics reported in the article considered only fully flanked hotspots (all islands located at the end of non-circular chromosomes were not considered).

The terminator sequences across all phage genomes were predicted using BATTER (Jin *et al*., 2023). Origin of transfer for plasmid sequences were predicted using oriTfinder (Li *et al*, 2018). DGR VR and TR location was annotated using myDGR tool (Sharifi & Ye, 2019).

Additional domain annotation for selected proteins were performed using HHPred (Soding *et al*, 2005), their structure prediction with AlphaFold3 (Abramson *et al*., 2024), and searching for structural homologues with FoldSeek (van Kempen *et al*, 2024). Protein structures were visualised using ChimeraX (Meng *et al*, 2023) Statistical analysis of results was performed in R with ggplot library used for visualisation (Wickham, 2016). Sequence logos were plotted using ggseqlogo (Wagih). Locus visualisation was performed using LoVis4u tool (Egorov & Atkinson, 2024).

## Data availability

iLund4u source code is provided on the GitHub page (github.com/art-egorov/ilund4u). The precomputed phage and plasmid iLund4u databases are available on our server: data-sharing.atkinson-lab.com/iLund4u/ and downloadable using command-line interface of iLund4u using: “ilund4u --database phages” or “ilund4u --database plasmids” commands. Results folder of iLund4u annotation are also available on the server in the folders *iLund4u_Phages* and *iLund4u_Plasmids*, for phage and plasmid results, respectively.

## Acknowledgments

This work was supported by the Knut and Alice Wallenberg Foundation (project grant 2020-0037 to G.C.A. and V.H.), the Swedish Research Council (Vetenskapsrådet) grants (2021-01146 to V.H., and 2022-01603 and 2023-02353 to G.C.A.), Cancerfonden (20 0872 Pj to V.H.), Göran Gustafsson Foundation for Research in Natural Sciences and Medicine (the Göran Gustafsson Prize to V.H.) and eSSENSE (eSSENCE@LU 10:2): the e-Science collaboration to GCA. Structural analyses were enabled by the supercomputing resources Berzelius provided by the National Supercomputer Centre (NSC) at Linköping University and the Knut and Alice Wallenberg foundation. Additional computational resources were provided by the National Academic Infrastructure for Supercomputing in Sweden (NAISS) and the Swedish National Infrastructure for Computing at NSC, Chalmers University Centre for Computational Science and Engineering (C3SE), LUNARC, The Centre for Scientific and Technical Computing at Lund University, and PDC Centre for High Performance Computing, KTH Royal Institute of Technology, partially funded by the Swedish Research Council through grant 2022-06725.

## Notes

### Competing Interest Statement

The authors have declared no competing interest.

