## Supplementary Information for "Systematic annotation of hyper-variability hotspots in phage genomes and plasmids"

[#] To whom correspondence should be addressed:

#### Supplementary Figures

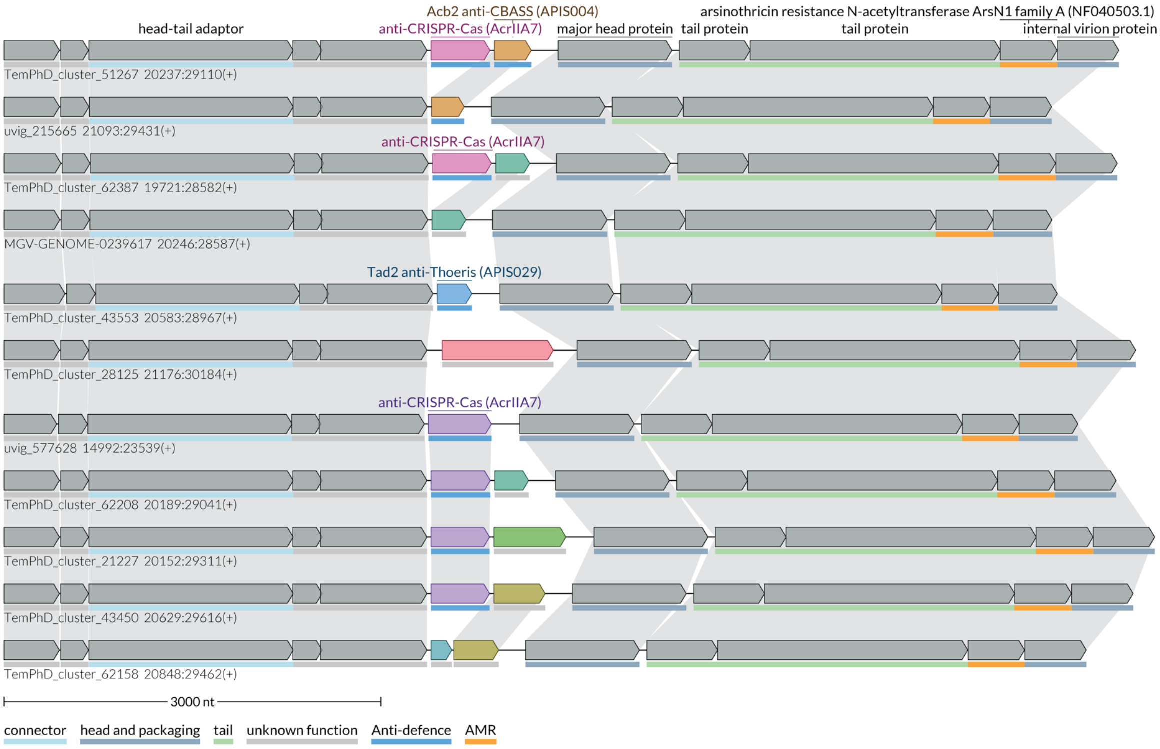


**Supplementary figure S1 | Anti-defence hotspot.**

Detailed view of the hotspot, located downstream of the head-tail adaptor and upstream of major head protein. Protein labels are hidden for hypothetical proteins or proteins of unknown function. The functions of annotated proteins are indicated by colored lines beneath each locus, with the color code provided at the bottom of the panel.

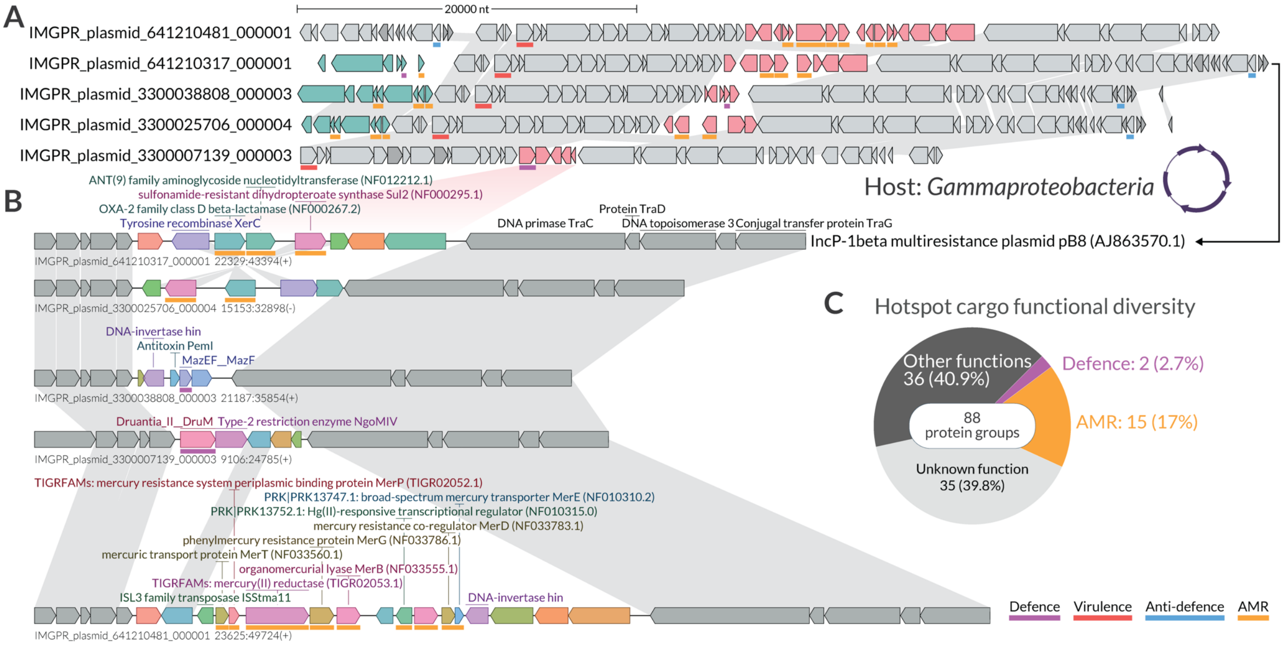


**Supplementary Figure S2 | Hotspot in the lagging region of the conjugative plasmids.** **(A)** LoVis4u visualization of a subset of plasmid sequences, highlighting its single hotspot where cargo proteins are shown in red, with all other proteins in grey. **(B)** Detailed view of the hotspot. Protein labels are hidden for hypothetical proteins or proteins of unknown function. The functions of annotated proteins are indicated by colored lines beneath each locus, with the color code provided at the bottom of the panel. **(C)** Pie charts of functional diversity of hotspot cargo protein groups.

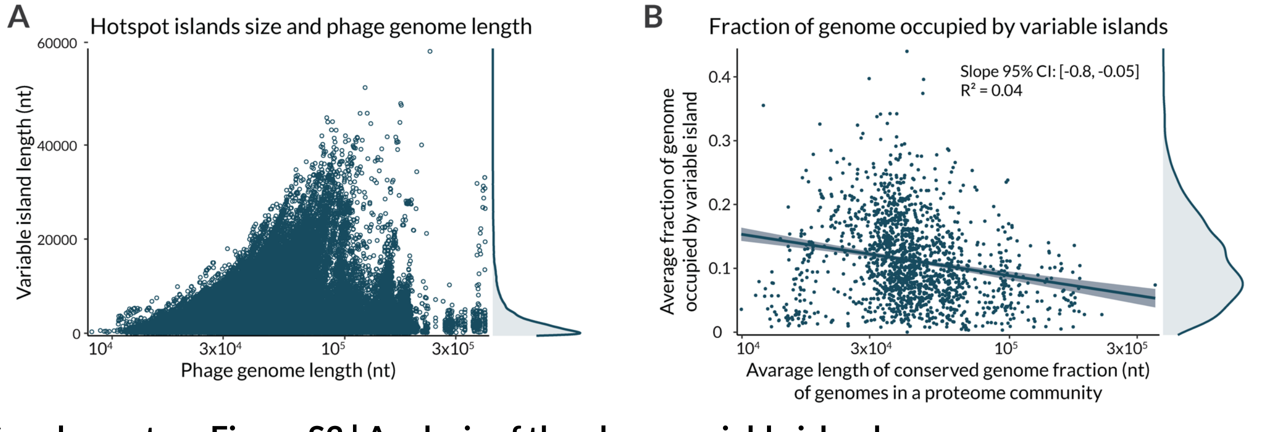


**Supplementary Figure S3 | Analysis of the phage variable islands.**

**(A)** Scatter plot of individual variable island sizes versus phage genome lengths (in nucleotides). **(B)** Correlation between the average fraction of the genome occupied by variable islands and the average length of the conserved genome fraction (in nucleotides).

### Supplementary Text S1

#### Algorithm description

##### Hotspot annotation mode

###### Defining groups of protein homologues and proteome filtering

The first step is reading the input data in GFF format with the initial filtering round based on proteome size. By default, proteomes with fewer than 15 proteins are excluded from the analysis. After GFF files are read, we use the MMseqs2 (Steinegger & Soding, 2017) clustering pipeline with all encoded proteins as input. We then treat each cluster of proteins as a group of homologues. The second filtering process at this step is exclusion of proteomes for which proteome composition uniqueness is less than a set cutoff, which by default is 0.7). Proteome composition uniqueness for $i^{th}$ proteome defined as $|G_i|/N_i$, where $|G_i|$ is the length (number of unique elements) of the set of protein groups and $N_i$ is the number of CDS. This is to exclude cases where large and potentially artefactual DNA duplications appear in the assembly.

###### Proteome network construction and community detection

The key step of the method is building the network of proteomes and defining communities of sequences that have similar proteome composition. In other words, we want to cluster input genomes based on fraction of shared homologues without using any information about taxonomy and nucleotide sequence alignments. In the next steps, this clusters will be used to define "*conserved*" and "*variable*" protein groups for each cluster that are equivalent to "*core*" and "*cloud*" terms of pan-genome analysis (Brockhurst *et al*, 2019).

The iLund4u proteome network is a weighted graph where each node, say $n_i$, represents the corresponding $i^{th}$ proteome. Each graph edge, say $e_{ij}$, connecting $i^{th}$ and $j^{th}$ proteomes, has weight $w_{ij}$:

$$w_{ij} = \left(\frac{|G_i \cap G_j|}{|G_i|} + \frac{|G_i \cap G_j|}{|G_j|}\right) \cdot \frac{1}{2} = \frac{|G_i \cap G_j| \cdot (|G_i| + |G_j|)}{2|G_i||G_j|};\ \ w_{ij} \in [0,1];\ \ w_{ij} = w_{ji}$$

where $|G_i|$ , $|G_j|$ are lengths (number of unique elements) of the sets of protein groups of $i^{th}$ and $j^{th}$ proteomes, respectively. $|G_i \cap G_j|$ is the number of shared (overlapped) protein groups between $i^{th}$ and $j^{th}$ proteomes. That is, $w_{ij}$ is a metric of similarity of two genomes in terms of their proteome composition.

While building the graph we write only edges where the weight passes the cutoff, which is by default $w_{ij} \geq 0.7$. In doing so we filter out connections with less similarity, and speed up the network construction, as well as reducing used disk space for network storage.

The complexity of the network constructions step is $O(N_p \cdot \overline{|G|} \cdot \bar{P})$, where $N_p$ is the number of proteomes, $\overline{|G|}$ is the average proteome size, and $\bar{P}$ is the average size of protein groups of homologues (in how many proteomes its members are encoded). In general, it equals the sum of shared homologues between proteomes that can be connected. In our tests, the elapsed times for the network building function on different architecture are as follows: i) 70 seconds for 20K Enterobacteria phage proteomes on M1 MacBook laptop; ii) 60 minutes for 563K phage proteomes on AMD 7413 (2.65 Ghz) computing node. We can compare it with PhamClust tool mentioned above, which requires 2.5 hours of runtime on a laptop for 5K phage genomes (Gauthier & Hatfull, 2023).

Once the network is constructed, the next step is to define communities of proteomes to reveal the substructure of more and less closely related proteomes within the network. For this task we use the Leiden algorithm (Traag *et al*, 2019) with the quality function based on the Constant Potts Model (CPM) (Traag *et al*, 2011):

$$Q = \sum_c \left[ e_c - \gamma \binom{n_c}{2} \right]$$

where $e_c$ is the weighted number of edges inside community $c$, $n_c$ is the number of nodes in community $c$, and $\gamma$ is the resolution parameter, which can be interpreted as the inner and outer edge density threshold and set as $\gamma = 0.5$ by default.

**Defining protein group classes**

Within each proteome community we can find homologous protein groups that are encoded in different fractions of the proteomes. Those that are found in the majority of proteomes we call *"conserved"* (equivalent to "core" in pan-genome analyses). Those protein groups that are found in a relatively small fraction of proteomes then we call *"variable"* (equivalent to "cloud" in pan-genome analyses). Finally, those protein groups that are neither very variable nor very conserved, we refer to as *"intermediate"* ("shell" in pan-genome analyses). Default cutoffs are $> 0.75$ for being *"conserved"* and $< 0.25$ for being *"variable"*.

**Annotation of variable islands**

After assigning the protein group class to each protein within each proteome community, we can now define *"variable island"* objects. We say that a *"variable island"* is a region containing a set of adjacent and non-conserved proteins with at least one protein being "variable". In other words, a *"variable island"* is a locus of "variable" protein(s) without internal interruption by "conserved" ORFs but possibly containing "intermediate" proteins. Here it is important to note that the annotation process also depends on the "circularity" property of a genome. iLund4u allows setting this property independently for each genome.

The most important island attribute that defines all subsequent steps is a set of its conserved flanking protein groups. When a *variable island* region is defined, we iterate over its flanking proteins encoded on both sides. Over iteration we add each "conserved" protein to the attribute of conserved neighbours (either left or right). The iteration on each side stops either when five neighbours are collected on each side or when we pass over eight neighbour proteins. The minimal number of collected "conserved" neighbours is set by default as five and if an island does not reach the total number of five "conserved" neighbours it will not be taken in subsequent steps.

**Island network construction and hotspot annotation**

We define a hotspot as a set of *variable islands* encoded in different genomes, but in the same locus. That is, they have similar "conserved" flanking genes. From this definition we can consider each variable island as a locus characterised by its “conserved” neighbours (more precisely by the set of protein groups of its “conserved” neighbours) and build the network of islands identical to the network of proteomes if we consider each island as a new proteome with its conserved flanking genes (without cargo proteins) as its proteins. That allows us to apply the same algorithm of network construction and community detection as already was described in the section of proteome network building.

When the network of islands is constructed in each proteome community, and we find communities in this network we can postulate that each community is a hotspot. From this definition we can see that a hotspot represents a cluster in a network with a high density of inner connections, which means they share a similar set of “conserved” flanking protein groups. At this step we add filtering. Firstly, the presence cutoff: a hotspot is kept only if it is found in at least of 0.3 of proteomes within a proteome community. In other words, clusters of variable islands that are found only in a small fraction of proteomes within a proteome community are not considered as hotspots. Additionally, if we have several islands on the same proteome clustered together in one hotspot then only the one with the highest sum of weights relative to other community members will be kept in the hotspot. This can happen if two islands are divided by few (one to three) conserved proteins and this step allows us to keep only the one that has a higher overlapping rate of flanking protein groups compared with the others.

**Hotspot merging**

We can expect that hotspots found in different proteome communities can have the same conserved neighbours due to the modularity of mobile elements such as phages or plasmids (Dion *et al*, 2020; Hendrix *et al*, 1999; Pesesky *et al*, 2019). In order to detect and merge such hotspots together we build the network of hotspots and apply the Leiden algorithm for community detection in the same way as for proteome and island networks. In that case, for each hotspot we define the attribute “signature”, which represents a set of protein groups that are found in at least of 0.75 of its island's "conserved" flanking proteins. That is, hotspot signature is a set of protein groups that are found as “conserved” flanking genes in the majority of its islands. Then we can build a network of hotspots identical to a network of proteomes if we consider each hotspot as a new proteome with corresponding signature as its proteins.

**Additional functional annotation**

For additional annotation of proteins encoded either as cargo of annotated hotspot islands or as flanking genes we use pyhmmer (Eddy, 2011; Larralde & Zeller, 2023). In order to reduce the running time, we use representative proteins of each protein group as a query set for searching. As was defined above, each cluster of proteins is considered as a set of homologues, then, the search results are attributed to each protein group based on its representative sequence.

The list of used databases: AMRFinderPlus (AMR genes) (Feldgarden *et al*, 2021), DefenseFinder and CasFinder (Anti-phage defence genes) (Couvin *et al*, 2018; Tesson *et al*,

2024), Cas dbAPIS_Acr (Anti-defence) (Yan *et al*, 2024), VFDB (Virulence factors) (Liu *et al*, 2022).

##### Protein search mode

Guilt by association, embedding or co-localization are widely used for annotating proteins with unknown functions. Protein search mode finds all hotspots where homologues of a query protein are encoded in an iLund4u database of hotspots. The location, other island cargo genes, as well as a hotspot property could help a user to build a hypothesis about the protein functions of query proteins as well as to rank them for verification.

##### Proteome annotation mode

This mode is designed to annotate hotspots even if the query is a single proteome, but it has a community of similar proteomes in the iLund4u database. Additionally, it gives the ability to define "conserved" proteins in your query proteome, whose homologues are found in the majority of similar proteomes and variable islands.
